## Supplementary File 1 for "Neural circuit mechanisms for transforming learned olfactory valences into wind-oriented movement"

| **Key Resources Table** | | | | |
| --- | --- | --- | --- | --- |
| **Reagent type (species) or resource** | **Designation** | **Source or reference** | **Identifiers** | **Additional information** |
| strain, strain background (Drosophila melanogaster) | *20xUAS-CsChrimson-mVenus attP18* | Klapoetke et al., 2014; PMID: 24509633 | N.A. |  |
| strain, strain background (Drosophila melanogaster) | *10XUAS-Chrimson88-tdTomato attP1* | Klapoetke et al., 2014; PMID: 24509633 | N.A. |  |
| strain, strain background (Drosophila melanogaster) | 13XLexAop2-IVS-ChrimsonR-mVenus-p10 attP18 | Vivek Jayaraman | N.A. |  |
| strain, strain background (Drosophila melanogaster) | 20XUAS-syn21-mScarlet-opt-p10 su(Hw)attp8 | Glenn Turner | N.A. |  |
| strain, strain background (Drosophila melanogaster) | *pJFRC200-10xUAS-IVS-myr::smGFP-HA in attP18* | Nern et al.,2015; PMID: 25964354 | N.A. |  |
| strain, strain background (Drosophila melanogaster) | *pJFRC225-5xUAS-IVS-myr::smGFP-FLAG in VK00005* | Nern et al.,2015; PMID: 25964354 | N.A. |  |
| strain, strain background (Drosophila melanogaster) | *pBPhsFlp2::PEST in attP3* | Nern et al.,2015; PMID: 25964354 | N.A. |  |
| strain, strain background (Drosophila melanogaster) | *pJFRC201-10XUAS-FRT>STOP>FRT-myr::smGFP-HA in VK0005* | Nern et al.,2015; PMID: 25964354 | N.A. |  |
| strain, strain background (Drosophila melanogaster) | *pJFRC240-10XUAS-FRT>STOP>FRT-myr::smGFP-V5-THS-10XUAS-FRT>STOP>FRT-myr::smGFP-FLAG_in_su(Hw)attP1* | Nern et al.,2015; PMID: 25964354 | N.A. |  |
| strain, strain background (Drosophila melanogaster) | MB043-split-LexA | This paper | N.A. | Available from Aso lab |
| strain, strain background (Drosophila melanogaster) | *empty-split-GAL4 (p65ADZp attP40, ZpGAL4DBD attP2)* | Seeds et al., 2014; PMID: 25139955 | N.A. |  |
| strain, strain background (Drosophila melanogaster) | MB001B (R10H10-p65ADZp in attP40 and R55A07ZpGAL4DBD in attP2) | This paper | N.A. | Available from Aso lab |
| strain, strain background (Drosophila melanogaster) | MB002B (R12C11-p65ADZp in attP40 and R14C08ZpGAL4DBD in attP2) | Aso 2014 eLife, DOI: 10.7554/eLife.04577 | N.A. |  |
| strain, strain background (Drosophila melanogaster) | MB011C (R14C08-p65ADZp in VK00027 and R15B01ZpGAL4DBD in attP2) | Aso 2014 eLife, DOI: 10.7554/eLife.04577 | N.A. |  |
| strain, strain background (Drosophila melanogaster) | MB018B (R20G03-p65ADZp in attP40 and R19F09ZpGAL4DBD in attP2) | Aso 2014 eLife, DOI: 10.7554/eLife.04577 | N.A. |  |
| strain, strain background (Drosophila melanogaster) | MB022B (R24E06-p65ADZp in attP40 and Tdc2ZpGAL4DBD in attP2) | Aso 2014 eLife, DOI: 10.7554/eLife.04577 | N.A. |  |
| strain, strain background (Drosophila melanogaster) | MB025B (R24E12-p65ADZp in attP40 and R52H01ZpGAL4DBD in attP2) | Aso 2014 eLife, DOI: 10.7554/eLife.04577 | N.A. |  |
| strain, strain background (Drosophila melanogaster) | MB027B (R24H08-p65ADZp in attP40 and R53F03ZpGAL4DBD in attP2) | Aso 2014 eLife, DOI: 10.7554/eLife.04577 | N.A. |  |
| strain, strain background (Drosophila melanogaster) | MB029B (R30G08-p65ADZp in attP40 and R11A03ZpGAL4DBD in attP2) | This paper | N.A. | Available from Aso lab |
| strain, strain background (Drosophila melanogaster) | MB032B (R30G08-p65ADZp in attP40 and THZpGAL4DBD in attP2) | Aso 2014 eLife, DOI: 10.7554/eLife.04577 | N.A. |  |
| strain, strain background (Drosophila melanogaster) | LH2456 (R38D01-p65ADZp in attP40 and R77F05ZpGAL4DBD in attP2) | Dolan 2019 eLife, DOI: 10.7554/eLife.43079 | N.A. |  |
| strain, strain background (Drosophila melanogaster) | MB043C (R58E02-p65ADZp in VK00027 and R32D11ZpGAL4DBD in attP2) | Aso 2014 eLife, DOI: 10.7554/eLife.04577 | N.A. |  |
| strain, strain background (Drosophila melanogaster) | MB077B (R25D01-p65ADZp in attP40 and R19F09ZpGAL4DBD in attP2) | Aso 2014 eLife, DOI: 10.7554/eLife.04577 | N.A. |  |
| strain, strain background (Drosophila melanogaster) | MB080C (R33E02-p65ADZp in attP40 and R50A05ZpGAL4DBD in attP2) | Aso 2014 eLife, DOI: 10.7554/eLife.04577 | N.A. |  |
| strain, strain background (Drosophila melanogaster) | MB082C (R40B08-p65ADZp in VK00027 and R23C06ZpGAL4DBD in attP2) | Aso 2014 eLife, DOI: 10.7554/eLife.04577 | N.A. |  |
| strain, strain background (Drosophila melanogaster) | MB083C (R52G04-p65ADZp in VK00027 and R94B10ZpGAL4DBD in attP2) | Aso 2014 eLife, DOI: 10.7554/eLife.04577 | N.A. |  |
| strain, strain background (Drosophila melanogaster) | MB109B (R76F05-p65ADZp in attP40 and R23C12ZpGAL4DBD in attP2) | Aso 2014 eLife, DOI: 10.7554/eLife.04577 | N.A. |  |
| strain, strain background (Drosophila melanogaster) | MB112C (R93D10-p65ADZp in VK00027 and R13F04ZpGAL4DBD in attP2) | Aso 2014 eLife, DOI: 10.7554/eLife.04577 | N.A. |  |
| strain, strain background (Drosophila melanogaster) | MB213B (R76F05-p65ADZp in attP40 and R32G08ZpGAL4DBD in attP2) | Aso 2014 eLife, DOI: 10.7554/eLife.04577 | N.A. |  |
| strain, strain background (Drosophila melanogaster) | MB296B (R15B01-p65ADZp in attP40 and R26F01ZpGAL4DBD in attP2) | Aso 2014 eLife, DOI: 10.7554/eLife.04577 | N.A. |  |
| strain, strain background (Drosophila melanogaster) | MB310C (R52G04-p65ADZp in VK00027 and R17C11ZpGAL4DBD in attP2) | Aso 2014 eLife, DOI: 10.7554/eLife.04577 | N.A. |  |
| strain, strain background (Drosophila melanogaster) | MB315C (R58E02-p65ADZp in attP40 and R48H11ZpGAL4DBD in attP2) | Aso 2014 eLife, DOI: 10.7554/eLife.04577 | N.A. |  |
| strain, strain background (Drosophila melanogaster) | MB390B (R19B06-p65ADZp in attP40 and R59G08ZpGAL4DBD in attP2) | This paper | N.A. | Available from Aso lab |
| strain, strain background (Drosophila melanogaster) | MB433B (R30E08-p65ADZp in attP40 and R11C07ZpGAL4DBD in attP2) | Aso 2014 eLife, DOI: 10.7554/eLife.04577 | N.A. |  |
| strain, strain background (Drosophila melanogaster) | MB434B (R30E08-p65ADZp in attP40 and R53C10ZpGAL4DBD in attP2) | Aso 2014 eLife, DOI: 10.7554/eLife.04577 | N.A. |  |
| strain, strain background (Drosophila melanogaster) | MB441B (R30G08-p65ADZp in attP40 and R48B03ZpGAL4DBD in attP2) | Aso 2014 eLife, DOI: 10.7554/eLife.04577 | N.A. |  |
| strain, strain background (Drosophila melanogaster) | MB555B (R73F07-p65ADZp in attP40 and R12D12ZpGAL4DBD in attP2) | This paper | N.A. | Available from Aso lab |
| strain, strain background (Drosophila melanogaster) | MB630B (VT026773-p65ADZp in attP40 and R72B05ZpGAL4DBD in attP2) | Aso&Rubin 2016 eLife, DOI: 10.7554/eLife.16135 | N.A. |  |
| strain, strain background (Drosophila melanogaster) | SS00096 (R19G02-p65ADZp in attP40 and R70G12ZpGAL4DBD in attP2) | Turner-Evans 2017 Elife, DOI: 10.7554/eLife.23496 | N.A. |  |
| strain, strain background (Drosophila melanogaster) | SS00504 (R25C01-p65ADZp in attP40 and R54H01ZpGAL4DBD in attP2) | This paper | N.A. | Available from Aso lab |
| strain, strain background (Drosophila melanogaster) | SS00543 (R33E06-p65ADZp in VK00027 and R89G09ZpGAL4DBD in attP2) | This paper | N.A. | Available from Aso lab |
| strain, strain background (Drosophila melanogaster) | SS00550 (R37A12-p65ADZp in attP40 and R27H08ZpGAL4DBD in attP2) | This paper | N.A. | Available from Aso lab |
| strain, strain background (Drosophila melanogaster) | SS00561 (R38E07-p65ADZp in attP40 and R55B01ZpGAL4DBD in attP2) | This paper | N.A. | Available from Aso lab |
| strain, strain background (Drosophila melanogaster) | SS00581 (R48H12-p65ADZp in attP40 and R41B07ZpGAL4DBD in attP2) | This paper | N.A. | Available from Aso lab |
| strain, strain background (Drosophila melanogaster) | SS00623 (R76F12-p65ADZp in attP40 and R33E06ZpGAL4DBD in attP2) | This paper | N.A. | Available from Aso lab |
| strain, strain background (Drosophila melanogaster) | SS01126 (R14C08-p65ADZp in attP40 and VT 037491ZpGAL4DBD in attP2) | This paper | N.A. | Available from Aso lab |
| strain, strain background (Drosophila melanogaster) | SS01227 (R14C08-p65ADZp in attP40 and R11F12ZpGAL4DBD in attP2) | This paper | N.A. | Available from Aso lab |
| strain, strain background (Drosophila melanogaster) | SS01262 (VT 029592-p65ADZp in attP40 and R19B06ZpGAL4DBD in attP2) | This paper | N.A. | Available from Aso lab |
| strain, strain background (Drosophila melanogaster) | SS01319 (VT 029592-p65ADZp in attP40 and R33H11ZpGAL4DBD in attP2) | This paper | N.A. | Available from Aso lab |
| strain, strain background (Drosophila melanogaster) | SS01126 (R14C08-p65ADZp in attP40 and VT 037491ZpGAL4DBD in attP2) | This paper | N.A. | Available from Aso lab |
| strain, strain background (Drosophila melanogaster) | SS32189 (VT033047-p65ADZp in attP40 and R32D10ZpGAL4DBD in attP2) | This paper | N.A. | Available from Aso lab |
| strain, strain background (Drosophila melanogaster) | SS32218 (VT 040712-p65ADZp in attP40 and R13D05ZpGAL4DBD in attP2) | This paper | N.A. | Available from Aso lab |
| strain, strain background (Drosophila melanogaster) | SS32219 (VT 013618-p65ADZp in attP40 and VT 063636ZpGAL4DBD in attP2) | This paper | N.A. | Available from Aso lab |
| strain, strain background (Drosophila melanogaster) | SS32228 (VT 040004-p65ADZp in attP40 and VT 020600ZpGAL4DBD in attP2) | This paper | N.A. | Available from Aso lab |
| strain, strain background (Drosophila melanogaster) | SS32230 (VT 029362-p65ADZp in attP40 and VT013618ZpGAL4DBD in attP2) | This paper | N.A. | Available from Aso lab |
| strain, strain background (Drosophila melanogaster) | SS32244 (R13F04-p65ADZp in attP40 and R20H08ZpGAL4DBD in attP2) | This paper | N.A. | Available from Aso lab |
| strain, strain background (Drosophila melanogaster) | SS32254 (R26H05-p65ADZp in attP40 and VT 049923ZpGAL4DBD in attP2) | This paper | N.A. | Available from Aso lab |
| strain, strain background (Drosophila melanogaster) | SS33909 (VT 026342-p65ADZp in attP40 and VT 033912ZpGAL4DBD in attP2) | This paper | N.A. | Available from Aso lab |
| strain, strain background (Drosophila melanogaster) | SS33917 (VT 007746-p65ADZp in attP40 and R64A11ZpGAL4DBD in attP2) | Yamada 2022; doi: https://doi.org/10.1101/2022.03.30.486484 | N.A. | Available from Aso lab |
| strain, strain background (Drosophila melanogaster) | SS33918 (VT 007746-p65ADZp in attP40 and R66B12ZpGAL4DBD in attP2) | Yamada 2022; doi: https://doi.org/10.1101/2022.03.30.486484 | N.A. | Available from Aso lab |
| strain, strain background (Drosophila melanogaster) | SS39541 (R84C10-p65ADZp in attP40 and R23E10ZpGAL4DBD in attP2) | This paper | N.A. | Available from Aso lab |
| strain, strain background (Drosophila melanogaster) | SS40549 (R23E10-p65ADZp in JK73A and R84C10ZpGAL4DBD in attP2) | This paper | N.A. | Available from Aso lab |
| strain, strain background (Drosophila melanogaster) | SS41731 (R23E10-p65ADZp in JK22C and R84C10ZpGAL4DBD in attP2) | This paper | N.A. | Available from Aso lab |
| strain, strain background (Drosophila melanogaster) | SS45222 (VT018689-p65ADZp in attP40 and VT048933ZpGAL4DBD in attP2) | Yamada 2022; doi: https://doi.org/10.1101/2022.03.30.486484 | N.A. | Available from Aso lab |
| strain, strain background (Drosophila melanogaster) | SS45234 (VT 026646-p65ADZp in attP40 and VT 029309ZpGAL4DBD in attP2) | Yamada 2022; doi: https://doi.org/10.1101/2022.03.30.486484 | N.A. | Available from Aso lab |
| strain, strain background (Drosophila melanogaster) | SS48882 (R54H04-p65ADZp in attP40 and R26C06ZpGAL4DBD in attP2) | This paper | N.A. | Available from Aso lab |
| strain, strain background (Drosophila melanogaster) | SS48899 (R89B06-p65ADZp in attP40 and VT 063627ZpGAL4DBD in attP2) | This paper | N.A. | Available from Aso lab |
| strain, strain background (Drosophila melanogaster) | SS48900 (R91F05-p65ADZp in attP40 and VT 054914ZpGAL4DBD in attP2) | This paper | N.A. | Available from Aso lab |
| strain, strain background (Drosophila melanogaster) | SS49755 (R56B05-p65ADZp in attP40 and R84B09ZpGAL4DBD in attP2) | This paper | N.A. | Available from Aso lab |
| strain, strain background (Drosophila melanogaster) | SS49897 (R26C06-p65ADZp in VK00027 and R54H04ZpGAL4DBD in attP2) | This paper | N.A. | Available from Aso lab |
| strain, strain background (Drosophila melanogaster) | SS49899 (R26C06-p65ADZp in su(Hw)attP8 and R54H04ZpGAL4DBD in attP2) | This paper | N.A. | Available from Aso lab |
| strain, strain background (Drosophila melanogaster) | SS56699 (TH-p65ADZp in VK00027 and VT025720ZpGAL4DBD in attP2) | Hulse 2021; https://doi.org/10.7554/eLife.66039 | N.A. |  |
| strain, strain background (Drosophila melanogaster) | SS67221 (VT 026646-p65ADZp in attP40 and VT 019911ZpGAL4DBD in attP2) | Yamada 2022; doi: https://doi.org/10.1101/2022.03.30.486484 | N.A. | Available from Aso lab |
| strain, strain background (Drosophila melanogaster) | SS88953 (VT014604-p65ADZp in attP40 and VT063740ZpGAL4DBD in attP2) | This paper | N.A. | Available from Aso lab |
| strain, strain background (Drosophila melanogaster) | SS88997 (R25G01-p65ADZp in JK22C and R15D05ZpGAL4DBD in attP2) | This paper | N.A. | Available from Aso lab |
| strain, strain background (Drosophila melanogaster) | *pJFRC100-20XUAS-TTS-Shibire-ts1-p10 in VK00005* | Pfeiffer et al., 2012: https://doi.org/10.1073/pnas.120452010 | N.A. |  |
| strain, strain background (Drosophila melanogaster) | UAS-TeNT | Keller et al., 2002: PMID: 11810637 | N.A. |  |
| antibody | anti-GFP (rabbit polyclonal) | Invitrogen | A11122  RRID:AB_221569 | 1:1000 |
| antibody | anti-Brp  (mouse monoclonal) | *Developmental Studies Hybridoma Bank* | nc82  RRID:AB_2341866 | 1:30 |
| antibody | anti-HA-Tag  (mouse monoclonal) | Cell Signaling Technology | C29F4; #3724  RRID:AB_10693385 | 1:300 |
| antibody | anti-FLAG  (rat monoclonal | Novus Biologicals | NBP1-06712  RRID:AB_1625981 | 1:200 |
| antibody | anti-V5-TAG Dylight-549  (mouse monoclonal) | Bio-Rad | MCA2894D549GA  RRID:AB_10845946 | 1:500 |
| antibody | anti-mous IgG(H&L) AlexaFluor-568  (goat polyclonal) | Invitrogen | A11031  RRID:AB_144696 | 1:400 |
| antibody | anti-rabbit IgG(H&L) AlexaFluor-488  (goat polyclonal) | Invitrogen | A11034  RRID:AB_2576217 | 1:800 |
| antibody | anti-mouse IgG(H&L) AlexaFluor-488 conjugated  (donkey polyclonal) | Jackson Immuno Research Labs | 715-545-151  RRID:AB_2341099 | 1:400 |
| antibody | anti-rabbit IgG(H&L) AlexaFluor-594  (donkey polyclonal) | Jackson Immuno Research Labs | 711-585-152  RRID:AB_2340621 | 1:500 |
| antibody | anti-rat IgG(H&L) AlexaFluor-647  (donkey polyclonal) | Jackson Immuno Research Labs | 712-605-153  RRID:AB_2340694 | 1:300 |
| antibody | anti-Mouse IgG (H&L) ATTO 647N  (goat polyclonal) | ROCKLAND | 610-156-121  RRID:AB_10894200 | 1:100 |
| antibody | anti-rabbit IgG (H+L) Alexa Fluor 568  (goat polyclonal) | Invitrogen | A-11036  RRID:AB_10563566 | 1:1000 |
| chemical compound, drug | 4-Methylcyclohexanol | VWR | AAA16734-AD |  |
| chemical compound, drug | Pentyl acetate | Sigma-Aldrich | 109584 |  |
| chemical compound, drug | Ethyl lactate | Sigma-Aldrich | W244015 |  |
| chemical compound, drug | Paraffin oil | Sigma-Aldrich | 18512 |  |
| software, algorithm | ImageJ and Fiji | NIH  Schneider et al., 2012 | <https://imagej.nih.gov/ij/>  <http://fiji.sc/> |  |
| software, algorithm | MATLAB | MathWorks | <https://www.mathworks.com/> |  |
| software, algorithm | Adobe Illustrator CC | Adobe Systems | <https://www.adobe.com/products/illustrator.html> |  |
| software, algorithm | GraphPad Prism 9 | GraphPad Software | <https://www.graphpad.com/scientific-software/prism/> |  |
| software, algorithm | Python | Python Software Foundation | <https://www.python.org/> |  |
| software, algorithm | neuPrint | HHMI Janelia | <https://doi.org/10.25378/janelia.12818645.v1> |  |
| software, algorithm | Cytoscape | (Shannon et al., 2003) | https://cytoscape.org/ |  |
| software, algorithm | NeuTu | [Zhao et al., 2018](https://elifesciences.org/articles/62576#bib209) | https://github.com/janelia-flyem/NeuTu |  |
| software, algorithm | ScanImage | Vidrio Technologies | https://vidriotechnologies.com/ |  |
| software, algorithm | VVDveiwer | HHMI Janelia | <https://github.com/takashi310/VVD_Viewer> |  |
| other | Grade 3MM Chr Blotting Paper | Whatmann | 3030-335 | Used in glass vials with paraffin-oil diluted odours |
| other | mass flow controller | Alicat | MCW-200SCCM-D | Mass flow controller used for the olfactory arena |
